# An intracellular pathogen response pathway promotes proteostasis in *C. elegans*

**DOI:** 10.1101/145235

**Authors:** Kirthi C. Reddy, Tal Dror, Jessica N. Sowa, Johan Panek, Kevin Chen, Efrem S. Lim, David Wang, Emily R. Troemel

**Author notes:** Corresponding author: Emily Troemel, 9500 Gilman Dr #0349, La Jolla, CA 92093.

## Abstract

Maintenance of proteostasis is critical for organismal health. Here we describe a novel pathway that promotes proteostasis, identified through the analysis of *C. elegans* genes upregulated by intracellular infection. We named this distinct transcriptional signature the Intracellular Pathogen Response (IPR), and it includes upregulation of several predicted ubiquitin ligase complex components such as the cullin *cul-6*. Through a forward genetic screen we found *pals-22*, a gene of previously unknown function, to be a repressor of the *cul-6/*Cullin gene and other IPR gene expression. Interestingly, *pals-22* mutants have increased thermotolerance and reduced levels of stress-induced polyglutamine aggregates, likely due to upregulated IPR expression. We found the enhanced stress resistance of *pals-22* mutants to be dependent on *cul-6*, suggesting that *pals-22* mutants have increased activity of a CUL-6/Cullin-containing ubiquitin ligase complex*. pals-22* mutant phenotypes are distinct from the well-studied heat shock and insulin signaling pathways, indicating that the IPR is a novel pathway that protects animals from proteotoxic stress.

## Introduction

Maintenance of protein homeostasis, or proteostasis, is crucial for organismal health. The protein quality control pathways involved in maintaining the integrity of the proteome include those that monitor and regulate protein synthesis, folding, and modification as well as those that facilitate degradation of damaged or misfolded proteins (Amm et al., 2014; Wolff et al., 2014). Disruption of proteostasis can lead to the accumulation of protein aggregates, which are associated with aging and many human diseases such as Alzheimer’s disease, Parkinson’s disease, and Huntington’s disease (Hipp et al., 2014; Morimoto, 2008; Taylor and Dillin, 2011).

Conditions that cause an increase in protein misfolding such as heat shock trigger protective responses that increase capacity for protein folding as well as protein degradation (Richter et al., 2010). A major pathway known to promote proteostasis is the highly conserved Heat Shock Response pathway, in which the transcription factor HSF-1 mediates a protective response to heat stress through the induced expression of many genes including chaperones, which bind to and aid in refolding proteins (Vihervaara and Sistonen, 2014). In addition, overexpression of HSF-1 has been shown to increase lifespan (Hsu et al., 2003), indicating that HSF-1-mediated stress response pathways protect against both acute thermal stress and the chronic stresses associated with aging. In *C. elegans*, HSF-1-mediated increase in lifespan requires the forkhead transcription factor DAF-16 (Hsu et al., 2003), which promotes longevity and stress resistance and functions in the conserved Insulin/IGF-1 signaling pathway (Lin et al., 1997; McColl et al., 2010; Webb and Brunet, 2014). DAF-16 acts together with HSF-1 to regulate expression of a number of genes that protect against proteotoxic stressors such as heat shock (Hsu et al., 2003).

In addition to pathways that increase protein folding capacity, there are also pathways that protect against proteotoxic stress by increasing protein degradation capacity, either through increased ubiquitylation or increased proteasome capacity (Amm et al., 2014). One pathway known to increase proteasome capacity is the “bounce-back response” in which a block in proteasome activity leads to upregulation of proteasome subunit gene expression (Radhakrishnan et al., 2010). This compensatory response is mediated by the transcription factor SKN-1 in *C. elegans* and the related Nrf1 in mammals, which are key transcription factors that promote oxidative stress resistance and can regulate longevity (Li et al., 2011; Radhakrishnan et al., 2014; Radhakrishnan et al., 2010). Upstream of the proteasome, there are ubiquitin ligases that monitor for misfolded proteins and target them for ubiquitylation and subsequent degradation. For example, the ubiquitin ligase CHIP is a ‘co-chaperone’ that can both bind to and ubiquitylate misfolded proteins to target them for degradation (Edkins, 2015). Interestingly, the role of CHIP in longevity has recently been shown to include its ability to ubiquitylate and target for degradation the insulin receptor DAF-2 (Tawo et al., 2017), which acts upstream of the DAF-16 transcription factor, providing a link between stress and longevity. Although CHIP is conserved and important for maintaining proteostasis, several studies indicate that there are other quality control ubiquitin ligases yet to be identified (Min et al., 2008; Morishima et al., 2008).

Through analysis of the *C. elegans* host response to intracellular infection, we have identified a response pathway that enhances proteostasis capacity through novel factors, including the predicted ubiquitin ligase component CUL-6. In prior work we performed gene expression analyses of the *C. elegans* transcriptional response to infection by the intracellular pathogen *Nematocida parisii* (Bakowski et al., 2014). This transcriptional response includes upregulation of genes encoding cullin-RING ubiquitin ligase components in the Skp-cullin-F-box (SCF) family and a number of genes of unknown function. These transcriptional changes are strikingly similar to the *C. elegans* response to infection by the Orsay virus, another natural intracellular pathogen of *C. elegans*, but are distinct from the response to extracellular pathogen infection (Bakowski et al., 2014; Chen et al., 2017; Sarkies et al., 2013). We have therefore named this common transcriptional signature the Intracellular Pathogen Response (IPR). Interestingly, perturbation of the ubiquitin-proteasome system, a key component of protein quality control, was also found to increase expression of some IPR genes including SCF ligase components (Bakowski et al., 2014).

Here we use a forward genetic screen to identify the gene *pals-22* as a negative regulator of IPR gene expression. We show that *pals-22* mutants have constitutive expression of a set of IPR genes including ubiquitin ligase complex components, and also have increased thermotolerance and reduced levels of stress-induced polyglutamine aggregates. Intriguingly, the increased stress resistance of *pals-22* mutants is dependent on the SCF ligase component CUL-6, suggesting that *pals-22* mutants may have upregulated activity of a CUL-6-containing ubiquitin ligase complex that can protect against proteotoxic stressors. The IPR appears to be distinct from previously described stress response pathways, as *pals-22* mutant phenotypes are not dependent on *daf-16* or *hsf-1*, and *pals-22* mutants do not have increased lifespan, in contrast to the increased lifespan of other stress resistant animals including those that overexpress *daf-16* or *hsf-1* (Henderson and Johnson, 2001; Morley and Morimoto, 2004; Munoz, 2003). Altogether, this work describes a novel pathway for stress resistance and provides insight into how animals regulate proteostasis capacity.

## Results

### *pals-22* is a negative regulator of IPR gene expression

To investigate the transcriptional regulation of genes induced by intracellular infection, we performed a forward genetic screen for mutations that cause altered expression of a GFP reporter for *pals-5*, which is among the genes most highly induced by microsporidia infection. *pals-5* is a gene of unknown function that is a member of an expanded gene family in *C. elegans* (Figure S1) called ‘*pals*’ for <u>p</u>rotein containing <u>ALS</u>2CR12 domain (Leyva-Diaz et al., 2017). This domain is named for its identification in the human gene ALS2CR12, which is located in a region implicated in the human disease amyotrophic lateral sclerosis 2 (ALS2) (Hentati et al., 1994) though is not believed to be the causative gene underlying the ALS2 disease (Hadano et al., 2001). Interestingly, many of the *pals* genes are highly induced by both microsporidia and viral infection (Figure S1) (Bakowski et al., 2014; Chen et al., 2017; Sarkies et al., 2013), and while they previously had no known function, genes like *pals-5* provide robust read-outs for intracellular infection. Under basal conditions, animals carrying the *pals-5p::GFP* reporter have little or no visible GFP expression (Figure 1A), while intracellular infection or proteasomal inhibition lead to greatly increased GFP expression (Bakowski et al., 2014). In this screen, we examined F2 progeny of mutagenized animals and identified three alleles (*jy1, jy2*, and *jy3*) that cause a strong increase in expression of the *pals-5p::GFP* reporter in basal conditions (Figure 1A-D). All three mutant alleles are recessive and map to LGIII; they also all fail to complement each other, indicating that they are mutations in the same gene. We found that the three alleles have essentially identical phenotypes and therefore used only the *jy1* and *jy3* alleles for most of our further analyses.

**Figure 1.**
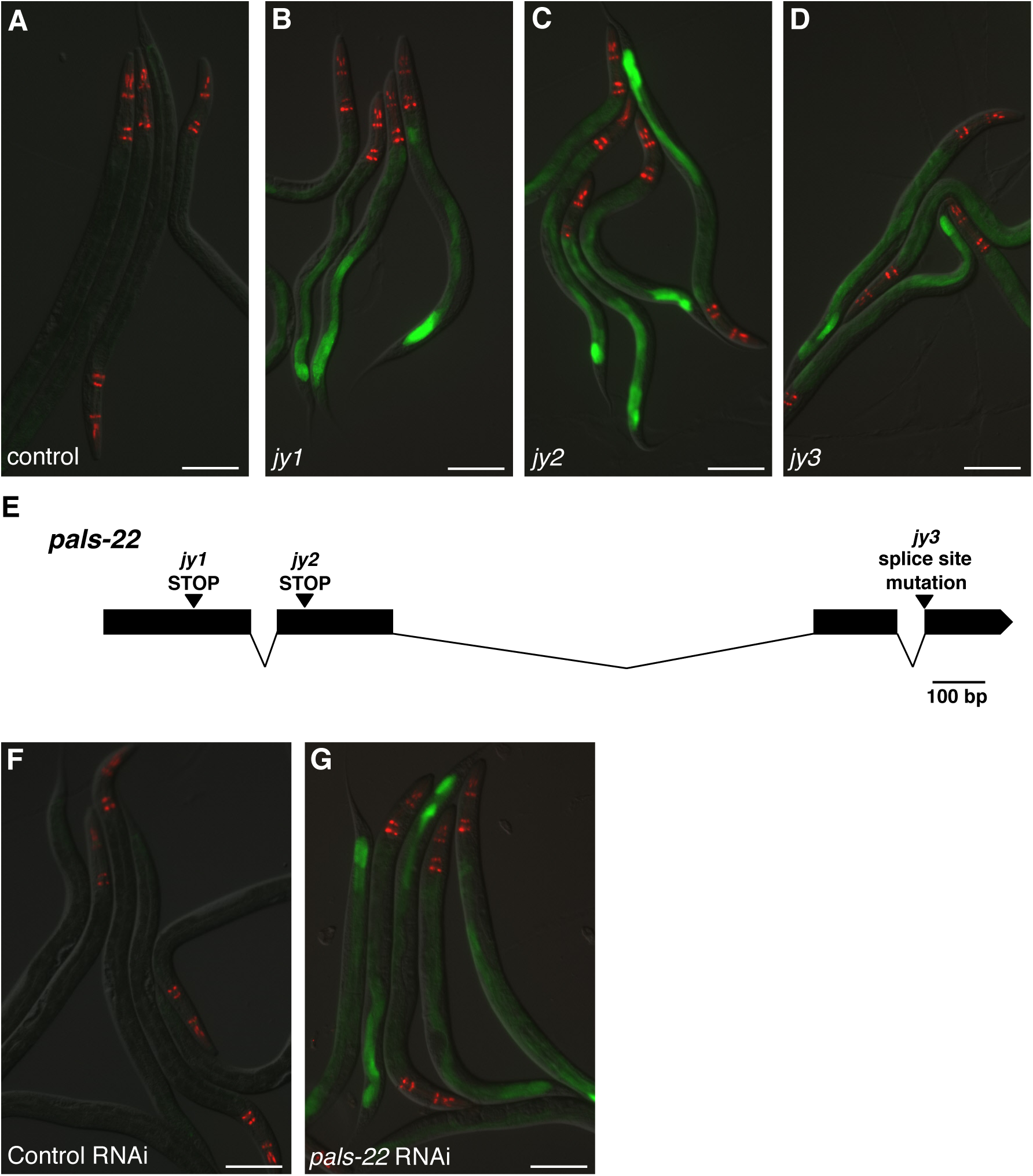
*pals-22* is a negative regulator of *pals-5p::GFP* expression. (A-D) Mutants isolated in a screen for increased expression of the *pals-5p::GFP* reporter. Shown are (A) control, (B) *pals-22(jy1)*, (C) *pals-22(jy2)*, and (D) *pals-22(jy3)* animals. (E) *jy1* and *jy2* encode stop mutations, and *jy3* alters a splice site in the *pals-22* gene. (F-G) *pals-5p::GFP* animals treated with either (F) L4440 RNAi control or (G) *pals-22* RNAi. In (A)-(D) and (F)-(G), green is *pals-5p::GFP*, red is *myo-2p::mCherry* expression in the pharynx as a marker for presence of the transgene. Images are overlays of green, red and Nomarski channels and were taken with the same camera exposure for all. Scale bar, 100 µm.

In addition to having altered *pals-5p::GFP* reporter expression, we observed that the *jy1* and *jy3* mutants develop more slowly than wild-type worms after hatching during normal well-fed conditions (Figure S2A-C). To quantify this effect, we measured the length of the *jy1* and *jy3* mutant worms at four different timepoints and found that the delay in growth is most pronounced after 48 hours of development (Figure S2E).

We used whole-genome sequencing analysis and identified the gene defective in *jy1, jy2*, and *jy3* mutants to be another member of the *pals* gene family, *pals-22*. Interestingly, the phylogenetic structure of the *pals* genes correlates with their regulation by infection, with *pals-22* in a clade of *pals* genes that are not induced by intracellular infection, in contrast to *pals-5*, which is in a clade that is induced by intracellular infection (Figure S1) (Bakowski et al., 2014; Chen et al., 2017; Sarkies et al., 2013). The predicted PALS-22 protein contains 284 amino acids with no predicted function or identifiable domains except the ALS2CR12 domain. The *jy1* and *jy2* alleles cause early stop mutations in *pals-22*, and *jy3* is a mutation predicted to affect splicing between exons 3 and 4 (Figure 1E). To confirm that *pals-22* is the gene responsible for the altered *pals-5* reporter expression in *jy1, jy2*, and *jy3* mutants, we performed RNAi of *pals-22* and found that it caused increased expression of the *pals-5p::GFP* reporter (Figure 1F-G), similar to the phenotypes of the *jy1, jy2*, and *jy3* mutants. In addition, we found that a fosmid containing the wild-type *pals-22* sequence tagged with GFP with native *cis*-regulatory elements (Sarov et al., 2012) could rescue the developmental delay of the *jy3* mutant (Figure S2D).

We next confirmed that these mutants have altered expression of endogenous *pals-5* mRNA expression by using qRT-PCR to measure RNA levels in whole animals in which the *pals-5p::GFP* reporter was not present. Indeed, we found that the *jy1* and *jy3* mutants have highly increased expression of *pals-5* mRNA as compared to wild-type animals (Figure 2), indicating that these *pals-22* mutations are not just affecting the *pals-5p::GFP* transgene. In addition, we measured levels of a subset of genes that are also induced as part of the IPR, including genes of unknown function *F26F2.1, F26F2.*3, and *F26F2.4* as well as the SCF ligase components *cul-6, skr-3, skr-4*, and *skr-5*. Interestingly, we found that all of the IPR genes in this subset have increased expression in the *jy1* and *jy3* mutants as compared to wild type, indicating that these mutants may have global defects in gene expression. We also used qRT-PCR to measure expression of another SCF component, *skr-1* (which is not induced by intracellular infection or proteasomal inhibition) (Bakowski et al., 2014), and found that this gene did not have increased expression in the *pals-22(jy1)* and *pals-22(jy3)* mutants (Figure 2). These results indicate that this subset of IPR genes are constitutively expressed in the *pals-22(jy1)* and *pals-22(jy3)* mutants, and that PALS-22 acts a negative regulator of transcription of these IPR genes.

**Figure 2.**
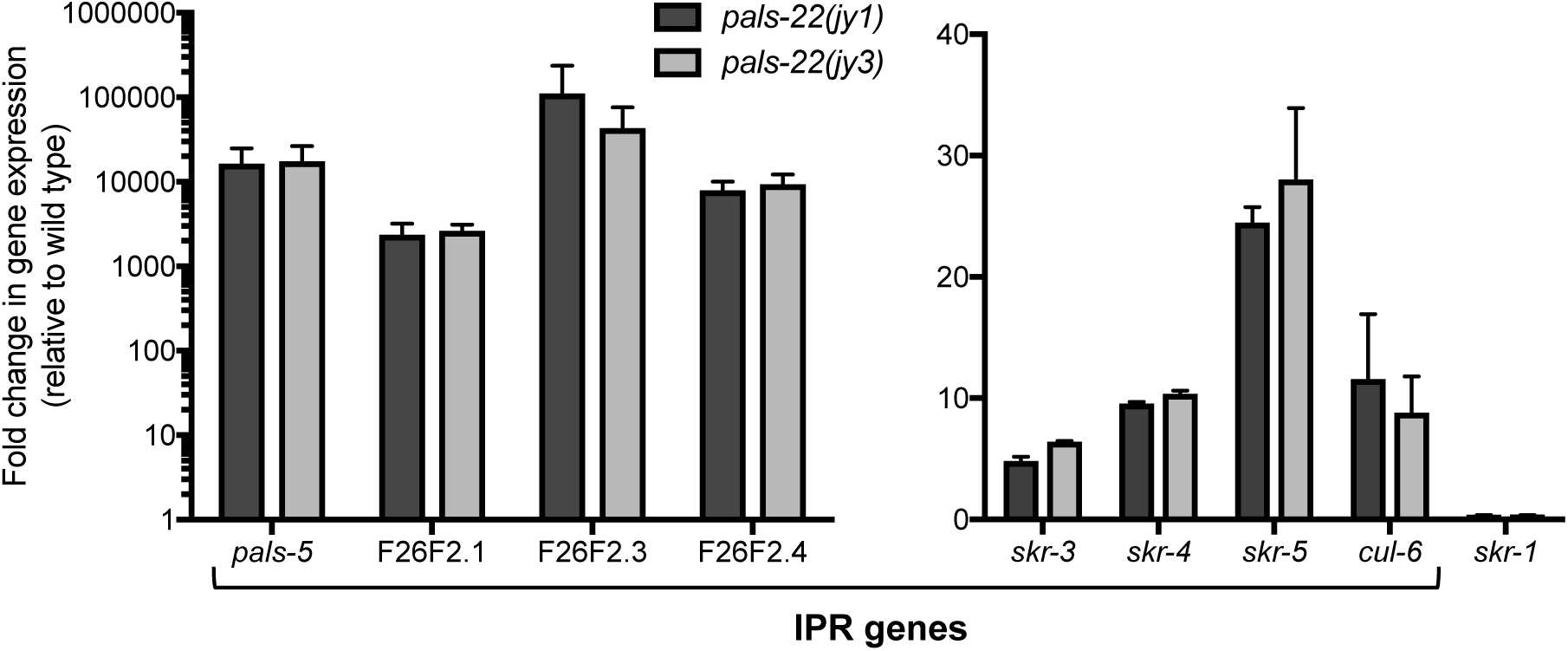
*pals-22* mutants have constitutive expression of IPR genes. qRT-PCR measurement of gene expression in *pals-22(jy1)* and *pals-22(jy3)* animals, shown as the fold change relative to wild-type control. Results shown are the average of two independent biological replicates, error bars are SD.

### *pals-22* mutants have increased thermotolerance, dependent on the cullin *cul-6*

It was previously reported that there is a partial overlap between the transcriptional response to intracellular infection and to a prolonged heat stress at 30°C, including genes in our IPR subset (*pals-5, F26F2.3, F26F2.4, skr-4*, and *skr-5*) (Bakowski et al., 2014; Mongkoldhumrongkul et al., 2010). We confirmed and extended this observation, finding that the *pals-5p::GFP* reporter has increased expression after prolonged heat stress (Figure S3A-B). We also used qRT-PCR to measure expression of genes in the IPR subset and found that these genes all had increased expression after prolonged heat stress treatment (Figure S3C).

This IPR induction in response to exposure to elevated temperature led us to test whether *pals-22* mutants have increased survival after heat shock treatment. In this assay we incubated worms at 37°C for 2 hours, and scored survival 24 hours later. We found that the *pals-22(jy1)* and *pals-22(jy3)* mutants had enhanced resistance to heat shock treatment compared to wild-type animals (Figure 3A), suggesting that constitutive IPR gene expression may have a protective effect against toxic effects of heat shock. We were able to rescue this phenotype using a transgene containing the wild-type *pals-22* gene, confirming that the increased thermotolerance phenotype results from loss-of-function of *pals-22* (Figure 3A).

**Figure 3.**
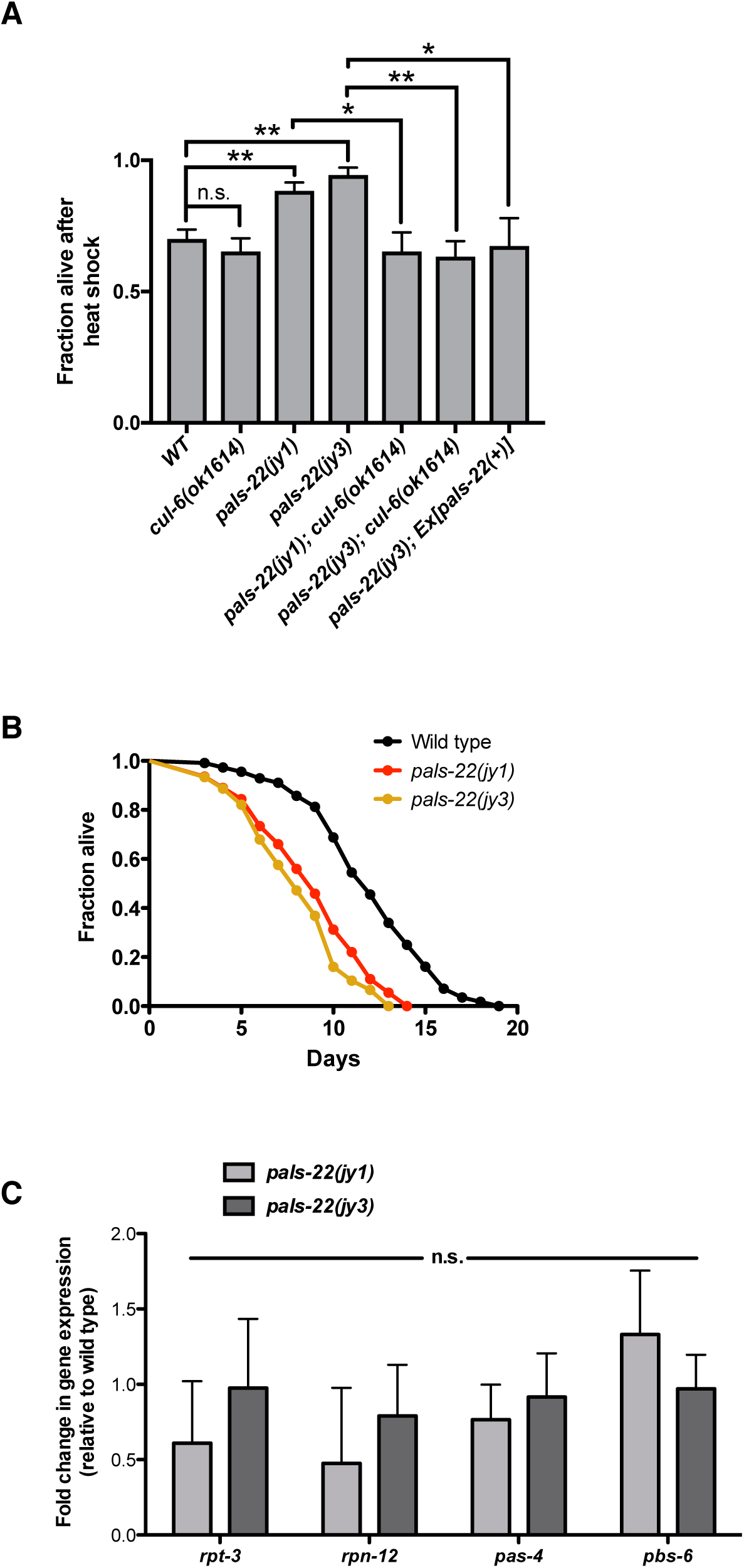
*pals-22* mutants have increased survival after heat shock, dependent on the cullin *cul-6*. (A) Survival of animals after 2 h heat shock treatment at 37°C. Strains were tested in triplicate. Mean fraction alive indicates the average survival among the triplicates, errors bars are SD. ** p < 0.01, * p < 0.05 with Student’s t-test. Heat shock assay was repeated three independent times with similar results, and data from a representative experiment are shown. (B) Lifespan of wild type, *pals-22(jy1)*, and *pals-22(jy3)* animals. Strains were tested in triplicate. Experiment was repeated two independent times with similar results, with data from a representative experiment shown. *p*-value for *pals-22(jy1)* and *pals-22(jy3)* compared to wild type is <0.0001 using the Log-rank test. (C) qRT-PCR measurement of proteasome subunit gene expression in *pals-22(jy1)* and *pals-22(jy3)* animals, shown as the fold change relative to wild-type control. Results shown are the average of two independent biological replicates, error bars are SD. n.s., not significant with Student’s t-test.

We next investigated whether the *pals-22* thermotolerance phenotype was dependent on any of the IPR genes. Mutation of the Skp1 homolog *skr-4* in a *pals-22* mutant background did not alter survival after heat shock (Figure S3D). Similarly, we also tested a large deletion encompassing 11 related *pals* genes (see Experimental Procedures) and did not see effects on *pals-22* thermotolerance (Figure S3E). In contrast, introducing the deletion allele *cul-6(ok1614)* into the *pals-22* mutant background had a striking effect. For both *pals-22* mutant alleles tested, the *cul-6* deletion reduced heat shock survival to levels comparable to wild type (Figure 3A), indicating that the increased thermotolerance of *pals-22* mutants is dependent on CUL-6 function. We introduced a fosmid containing the wild-type *cul-6* sequence with native *cis*-regulatory elements (Sarov et al., 2012) into the *pals-22(jy1);cul-6(ok1614)* double mutant to test for rescue but could not assay heat shock tolerance due to the sickness and decreased viability of this strain. Worms carrying the *cul-6(ok1614)* mutation alone had no reduction of thermotolerance as compared to wild type in our assays (Figure 3A), indicating that this gene does not significantly affect heat shock resistance in a wild-type context.

To address the possibility that the suppression of *pals-22* thermotolerance by *cul-6* was through effects on gene expression, we used qRT-PCR to measure mRNA levels of a set of IPR genes in *pals-22;cul-6* double mutants. We found that mutation of *cul-6* did not alter the constitutive expression of any of the genes tested (Figure S4A). We also tested whether the *cul-6* deletion suppressed the developmental delay of *pals-22* mutants, and did not see any effects on this phenotype (Figure S4B). Taken together, these results suggest that increased activity of CUL-6 in *pals-22* mutants promotes thermotolerance.

### *pals-22* phenotypes are independent of known stress response pathways

The transcription factors *hsf-1* and *daf-16* play important roles in the response to stress, including regulation of heat shock protein expression. We therefore investigated whether the *pals-22* constitutive IPR gene expression and increased thermotolerance phenotypes were independent of these well-characterized pathways. We generated the double mutant *hsf-1(sy441); pals-22(jy1)* but as this strain has a larval arrest phenotype we could not test effects of this *hsf-1* mutation on *pals-22* phenotypes. Due to technical difficulties, we were not able to test *hsf-1* RNAi in the heat shock assay (see Experimental Procedures). However, using RNAi we found that *hsf-1* is not required for the constitutive IPR gene expression phenotype of *pals-22* mutants (S5A-B).

Previous work indicated that the genes upregulated by intracellular infection are not regulated by *daf-16* (Bakowski et al., 2014). Importantly, we found that mutation of *daf-16* did not affect the *pals-22* increased thermotolerance phenotype (Figure S5C), supporting the model that stress resistance of *pals-22* mutants is independent of the canonical heat shock/insulin-signaling pathway involving *hsf-1* and *daf-16*.

Many manipulations in *C. elegans* that increase resistance to heat shock also confer a prolonged lifespan (Munoz, 2003). We therefore analyzed the lifespan of *pals-22(jy1)* and *pals-22(jy3)* mutant worms at 25°C. Surprisingly, we found that these strains have a decreased lifespan as compared to wild type (Fig 3B). This result suggests that while constitutive IPR gene expression may lead to stress resistance in younger animals, it may have deleterious effects on the health of the animal that result in early death.

Previous work has demonstrated that inhibition of proteasome function results in transcriptional upregulation of proteasome subunits, as part of a conserved “bounce-back” response (Li et al., 2011). To test whether inhibition of *pals-22* function leads to a similar response, we used qRT-PCR to examine expression of proteasome genes (Figure 3C). We found no increase in proteasome subunit mRNA levels in *pals-22* mutants as compared to wild type. Together these results suggest that increased stress resistance of *pals-22* mutants is distinct from previously described stress response pathways.

### *pals-22* mutants have decreased polyglutamine aggregation, dependent on *cul-6*

Given the increased stress resistance observed in *pals-22* mutants, we tested for another phenotype commonly found in stress-resistant animals. Specifically, we measured aggregation of a polyglutamine (polyQ) repeat – YFP fusion protein, which is derived from the polyQ repeats found on the huntingtin protein responsible for Huntington’s disease. We used an integrated transgene that expresses a polyQ(44)::YFP protein from the intestine-specific *vha-6* promoter (Mazzeo et al., 2012), and examined polyQ aggregate formation in *pals-22* mutants upon stress. It was previously reported that osmotic stress rapidly induces the formation of intestinal polyQ aggregates, which can be scored in live animals (Mazzeo et al., 2012). We found that *pals-22* mutants have a greatly reduced number of polyQ aggregates compared to control after osmotic stress treatment (Figure 4A-B). We monitored aggregates over a 24 hour period and observed that *pals-22* mutants have delayed aggregate formation – at early timepoints the *pals-22* mutants have reduced numbers of aggregates, but at the 24 hour timepoint aggregate formation in *pals-22* mutants is comparable to control worms (Figure 4C).

**Figure 4.**
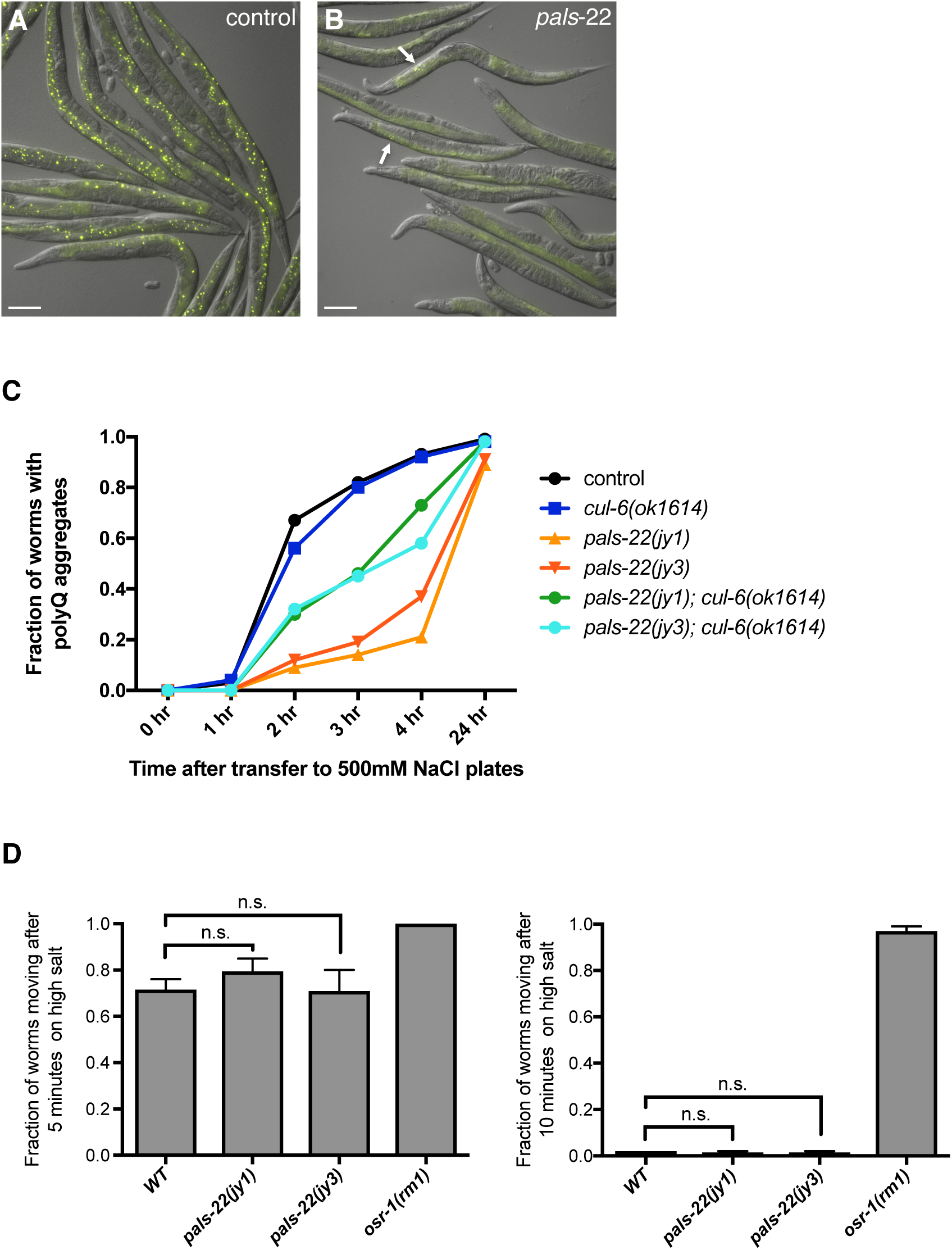
*pals-22* mutants have decreased polyQ aggregation, dependent on *cul-6*. (A,B) Images of (A) control and (B) *pals-22(jy3)* animals expressing polyQ(44)::YFP at 3 hours after transfer to high salt plates. Arrows indicate osmotic stress-induced aggregates. Images are overlays of yellow and Nomarski channels and were taken with the same camera exposure. Scale bar, 100 µm. (C) Fraction of animals with polyQ aggregates after transfer to high salt plates. At least 60 worms were scored for each strain. Assay was repeated two independent times with similar results, and data from a representative experiment shown. (D) Fraction of animals scored as moving after transfer to high salt plates at the indicated timepoints. At least 50 worms were scored for each strain. Results shown are the average of two independent replicates. Error bars are SD. n.s., not significant with Student’s t-test.

As worms resistant to osmotic stress had greatly decreased formation of osmotically-induced intestinal polyQ aggregates (Mazzeo et al., 2012), we tested whether the reduced polyQ aggregation phenotype of *pals-22* mutants could simply be due to a general resistance to osmotic stress. We transferred worms to high-salt plates and tested movement in response to touch over a period of 10 minutes. While the osmotic stress-resistant mutant *osr-1* continued to move throughout this assay, *pals-22* mutants had salt-induced paralysis comparable to wild-type worms (Figure 4D). This indicates that *pals-22* mutants are not resistant to osmotic stress but rather have a specific reduction in polyQ aggregate formation.

We next tested if this reduced aggregate formation phenotype was dependent on *cul-6*, similar to the thermotolerance phenotype of *pals-22* mutants. We found that *pals-22;cul-6* double mutants had an intermediate aggregate formation phenotype (Figure 4C), suggesting that *cul-6* functions to reduce polyQ aggregation in *pals-22* mutants, although there are likely other factors contributing to this phenotype as well.

Finally, we quantified the amount of polyQ(44)::YFP protein expressed in the strains used in our analysis to ensure that the altered aggregate formation was not due to changes in polyQ protein levels. We used a COPAS Biosort machine to measure YFP fluorescence in these worms and found that mutation of *pals-22* or *cul-6* does not alter the amount of polyQ(44)::YFP fluorescence (Figure S6A). We also used western blot analysis and similarly found no change in the amounts of polyQ(44)::YFP protein in all six strains tested (Figure S6B). Thus, *pals-22* does not appear to affect the overall levels of polyQ(44)::YFP protein, just the propensity to form stress-induced aggregates.

### PALS-22 is broadly expressed and functions in the intestine to regulate IPR gene expression and thermotolerance

We used fosmids containing GFP-3xFLAG-tagged versions of *pals-22* and *cul-6* with native *cis*-regulatory elements (Sarov et al., 2012) to identify the tissues in which these genes are expressed. We generated worms carrying these transgenes, and observed PALS-22::GFP expression throughout the animal, including expression in the intestine, neurons, and hypodermis (Figure 5A). We found that CUL-6::GFP was not as broadly expressed, with GFP seen only in the pharynx and intestine (Figure 5B).

**Figure 5.**
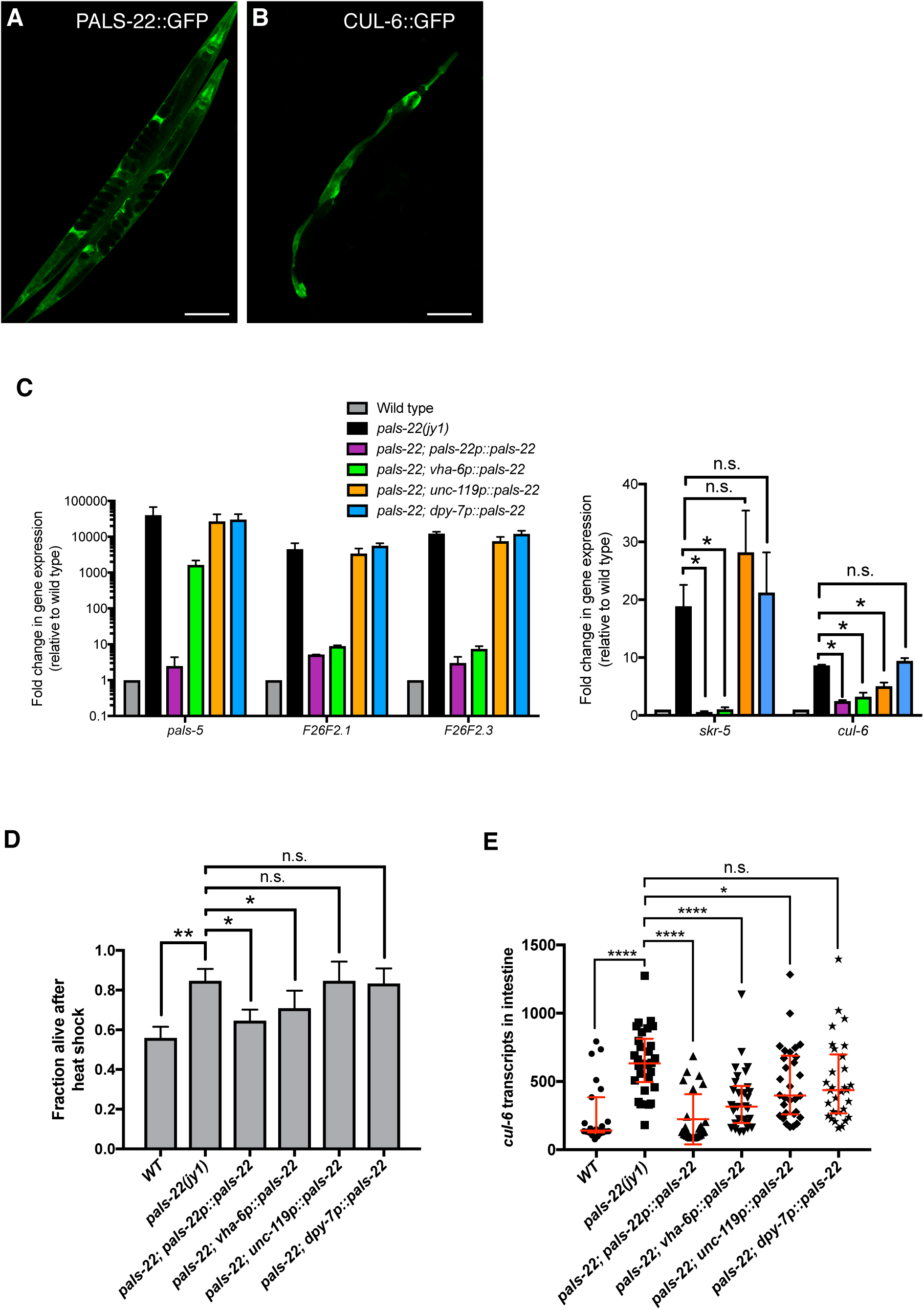
PALS-22 is broadly expressed, and functions in the intestine to regulate IPR expression and thermotolerance. (A,B) Confocal fluorescence images of adult animals with fosmid transgenes expressing (A) PALS-22::GFP and (B) CUL-6::GFP from endogenous promoters. (C) qRT-PCR measurement of IPR gene expression in animals with tissue-specific expression of *pals-22(+)* driven by endogenous (*pals-22*), intestinal (*vha-6*), neuronal (*unc-119*), or hypodermal (*dpy-7*) promoters in a *pals-22(jy1)* mutant background. Results are shown as the fold change relative to wild-type control and are the average of two independent biological replicates. Error bars are SD. * p < 0.05, n.s., not significant with Student’s t-test. (D) Survival of animals with *pals-22* tissue-specific rescue after 2 h heat shock treatment at 37°C. Strains were tested in triplicate. Mean fraction alive indicates the average survival among the triplicates, errors bars are SD. ** p < 0.01, * p < 0.05, n.s., not significant with Student’s t-test. Heat shock assay was repeated three independent times with similar results, and data from a representative experiment are shown. (E) Measurement of intestinal *cul-6* mRNA transcripts using single molecule fluorescence *in situ* hybridization (smFISH). Graph is a compilation of three independent biological replicates. Each symbol represents the smFISH spot count from the four anterior-most intestinal cells of an individual worm. Statistical analysis was performed using one-way ANOVA.

We next investigated in which tissues the PALS-22 protein acts to regulate gene expression and thermotolerance. We expressed the *pals-22* cDNA tagged with GFP using the endogenous *pals-22* promoter as well as intestinal, neuronal, and hypodermal promoters for tissue-specific expression. We generated single-copy insertion of these transgenes and crossed them into the *pals-22(jy1)* mutant background. Expression of the *pals-22* cDNA from the endogenous promoter rescued the constitutive IPR gene expression phenotype, as measured by qRT-PCR (Figure 5C) as well as the increased thermotolerance phenotype (Figure 5D), indicating that this tagged cDNA is functional. We found that intestinal expression of *pals-22* could strongly rescue the constitutive IPR gene expression phenotype of most genes tested, while pan-neuronal and hypodermal expression of *pals-22* did not result in large effects on IPR gene expression (Figure 5C). Similarly, we saw a partial rescue of the *pals-22* increased thermotolerance phenotype with intestinal rescue but no significant effect with neuronal or hypodermal rescue of *pals-22* (Figure 5D).

We used single molecule fluorescent in situ hybridization (smFISH) to quantify mRNA expression of *cul-6*, which is required for the *pals-22* mutant enhanced proteostasis phenotypes. In *pals-22(jy1)* worms, we observed increased expression of *cul-6* mRNA in the intestine as compared to wild-type worms, which was rescued by expression of *pals-22* from the endogenous promoter (Figure 5E). Expression of *pals-22* from either intestinal or neuronal promoters decreased the amount of *cul-6* mRNA seen in the intestine, with the intestinal *pals-22* expression having a larger effect. Similar to the thermotolerance and IPR gene expression phenotypes tested, we did not see a significant effect on intestinal *cul-6* expression with hypodermal expression of *pals-22* (Fig 5E). Taken together our results suggest that *pals-22* acting in the intestine regulates *cul-6* expression in this tissue, which may contribute to the increased thermotolerance of *pals-22* mutants.

## Discussion

In response to diverse kinds of natural intracellular infection, *C. elegans* induces a distinct transcriptional signature we are calling the Intracellular Pathogen Response (IPR). The IPR is characterized by transcriptional upregulation of genes of unknown function, such as genes in the large *pals* gene family (Figure S1), as well as genes that are predicted to encode ubiquitin ligase components in the SCF family, such as *cul-6*. Through a forward genetic screen for constitutive expression of the IPR gene *pals-5*, we identified another *pals* gene called *pals-22* that acts as a repressor of expression of a subset of IPR genes including *cul-6* and *pals-5.* Interestingly, a subset of IPR genes can be induced not just by intracellular infection, but also by proteotoxic stressors such as proteasome inhibition and heat stress. These diverse stress conditions are all expected to negatively impact cellular proteostasis, suggesting that the IPR is induced as a response to proteotoxic stress, which led us to analyze *pals-22* proteostasis phenotypes. Indeed, we found that *pals-22* mutants, which constitutively express a subset of IPR genes including *cul-6*, have increased resistance in thermotolerance and polyglutamine aggregation stress assays. These findings suggest that the IPR can improve proteostasis capacity in acute stress conditions (Figure 6).

**Figure 6.**
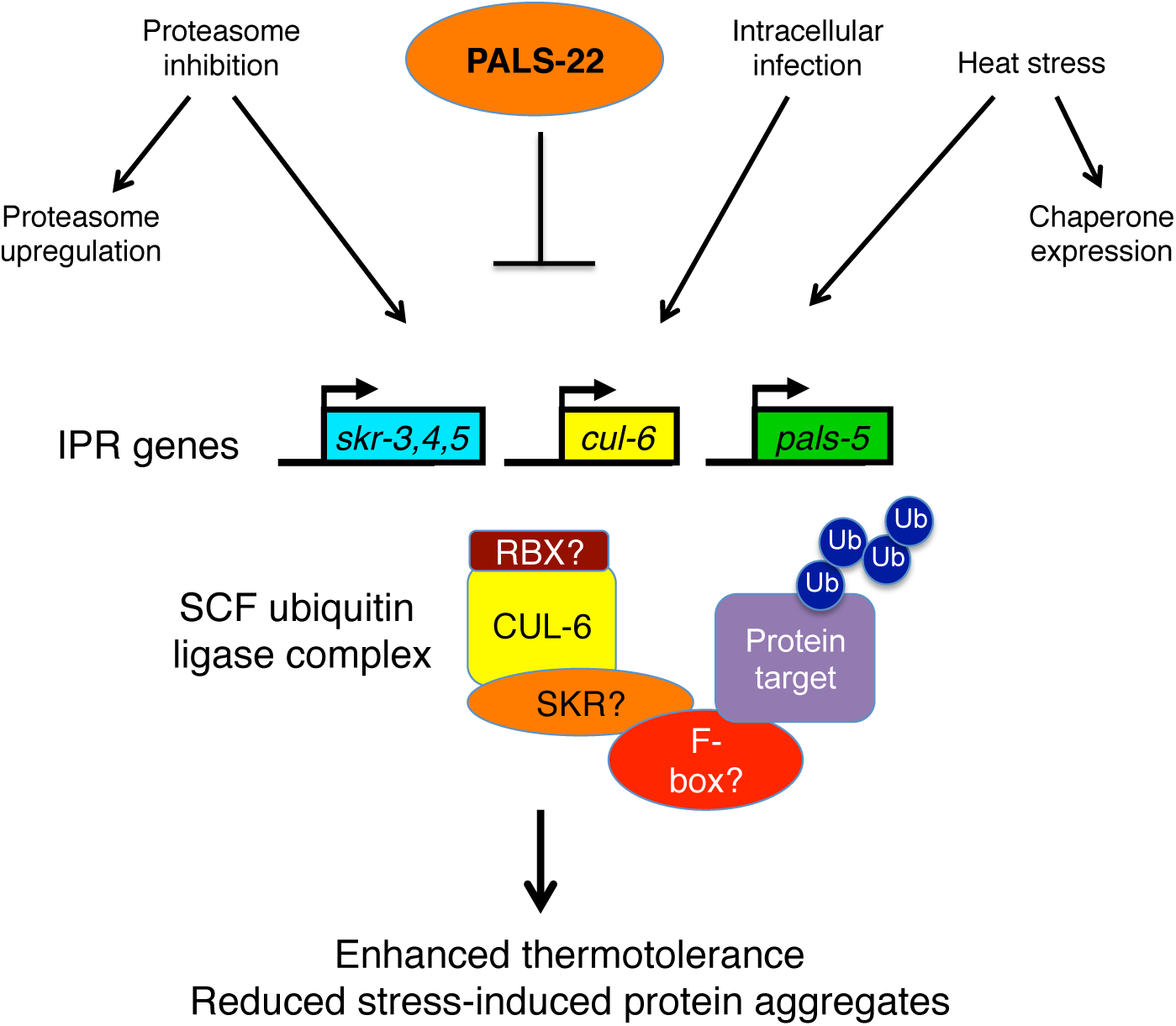
Model for increased proteostasis capacity in *pals-22* mutants.

The IPR appears to be distinct from previously described stress response pathways. The transcriptional response to intracellular infection does not include canonical heat shock proteins and does not have significant overlap with the HSF-1-regulated Heat Shock Response (HSR) or the Insulin/IGF-1 signaling pathway (Bakowski et al., 2014). In addition, our data indicate that *pals-22* mutant phenotypes are not dependent on *hsf-1* or *daf-16*. The IPR is also distinct from the previously reported “bounce-back” response, in which transcription of proteasome components is upregulated in response to proteasomal inhibition. These results suggest that the enhanced proteostasis capacity conferred by a constitutively activated IPR is not due to reduced proteasome substrate load (through upregulation of protein chaperones to cause refolding), or by increased capacity for proteasomal degradation (through upregulation of proteasome subunit genes), but rather through an increased targeting of misfolded proteins for degradation (through increased expression of ubiquitin ligase components). In contrast to mutants in these previously described proteostasis pathways, we found that *pals-22* mutants with constitutive IPR gene expression do not have extended lifespan, further supporting the hypothesis that *pals-22* regulates a distinct proteostasis pathway.

Notably, the increased stress resistance of *pals-22* mutants is dependent on the cullin *cul-6*. Cullins are known to act as core scaffolding components in multi-subunit cullin-RING ubiquitin ligases. *cul-6* is the only one of six cullins present in the *C. elegans* genome that is an IPR gene, and our previous work indicates that CUL-6 plays a role in the *C. elegans* defense against pathogen infection (Bakowski et al., 2014). We did not observe any decreased survival after heat shock or increased stress-induced polyQ aggregation in *cul-6* mutants as compared to wild type, indicating that while *cul-6* is critical for the increased proteostasis capacity of *pals-22* mutants, it is not normally needed for protection in these conditions. A previous study found that RNAi inhibition of the cullins *cul-1* or *cul-2* accelerated polyglutamine-induced paralysis in *C. elegans*, suggesting that these cullins may promote proteostasis in wild-type worms (Mehta et al., 2009).

Where are PALS-22 and CUL-6 acting to regulate proteostasis? We found that while PALS-22 is broadly expressed, it functions in the intestine to regulate thermotolerance. In addition, our results indicate that PALS-22 activity in the intestine regulates expression of *cul-6* mRNA in the intestine, suggesting that both PALS-22 and CUL-6 are acting in this tissue to regulate thermotolerance. The simplest model for how PALS-22 and CUL-6 interact in the intestine is that PALS-22 is a negative regulator of *cul-6* transcription. However, as neither our work nor a parallel study (Leyva-Diaz et al., 2017) found PALS-22 protein enriched in the nucleus, PALS-22 is likely not a transcription factor but rather may be an indirect regulator of *cul-6* mRNA expression. However, it is possible that PALS-22 also functions biochemically with CUL-6 and acts as a more direct inhibitor of a process mediated by a CUL-6-containing ubiquitin ligase in the intestine.

Regardless of the interaction between PALS-22 and CUL-6, CUL-6 is predicted to act as a component of a multi-subunit cullin-RING ubiquitin ligase complex. Increased expression of such a ubiquitin ligase complex in *pals-22* mutants could then lead to increased ubiquitylation activity and perhaps increased targeting of misfolded or toxic proteins to improve proteostasis (Figure 6). What other SCF ligase components could be acting with CUL-6? SCF ligase components canonically form a complex composed of a RING-containing RBX protein, SKP, cullin, and F-box-containing proteins (Hua and Vierstra, 2011). *C. elegans* has at least 21 SKP-related proteins, three of which are induced as part of the IPR (*skr-3, skr-4*, and *skr-5*). Previous work found that CUL-6 and SKR-3 can interact physically in a yeast two-hybrid assay, indicating these components could function together in an SCF ligase complex (Nayak et al., 2002). Mutation of *skr-4* did not affect the *pals-22* thermotolerance phenotype, but it is possible that one of the other SKP-related proteins acts together with CUL-6.

Related to our findings reported here, other studies have found that overexpression of ubiquitin ligases can result in increased stress resistance or increased lifespan. For example, in the yeast *Saccharomyces cerevisiae* it has been shown that overexpression of the ubiquitin ligase Rsp5 conferred resistance to various stressors, including heat stress (Hiraishi et al., 2006). Work done in plants has found that constitutive expression of ubiquitin ligases led to improved tolerance of high salt or drought conditions (Seo et al., 2012; Zhou et al., 2010). In *C. elegans*, overexpression of the HECT E3 ubiquitin ligase WWP-1 was shown to extend lifespan (Carrano et al., 2009).

Interestingly, a parallel study found that *pals-22* mutants have increased transgene silencing and an enhanced exogenous RNAi response, suggesting that the PALS-22 protein can regulate levels of small RNAs (Leyva-Diaz et al., 2017). These findings may be linked to our observation of increased gene expression in *pals-22* mutants. As *pals-22* mutants have an enhanced exogenous RNAi response it is possible they may also have a reduced endogenous RNAi response, as previous studies have shown exogenous and endogenous RNAi pathways are inversely correlated (Fischer et al., 2011). Therefore, IPR gene expression might normally be repressed by endogenous RNAi pathways, which are then derepressed in *pals-22* mutants leading to increased expression of IPR genes. Our future studies will focus on the mechanism used by *pals-22* to repress expression of IPR genes like *cul-6*, as well as the biochemical mechanism by which PALS-22 and CUL-6 increase proteostasis capacity to promote resistance to diverse stressors.

## Experimental Procedures

### Strains

*C. elegans* were maintained on NGM plates seeded with Streptomycin-resistant OP50-1 under standard conditions at 20°C as described (Brenner, 1974) unless otherwise mentioned. We used N2 wild-type animals. See Table 1 for list of all strains used in this study. Transgenic strains were generated as described previously (Frøkjær-Jensen et al., 2008; Kadandale et al., 2009). Transgenic strains expressing TransgeneOme fosmids (Sarov et al., 2012) as extrachromosomal arrays were generated by injecting fosmids at 100 ng/ul into strain EG6699 and selecting non-unc animals.

### EMS screen and cloning of alleles

Worms carrying the *jyIs8[pals-5p::GFP, myo-2::mCherry]* transgene were mutagenized with ethyl methane sulphonate (EMS) using standard procedures as described (Kutscher and Shaham, 2014), and screened in the F2 generation for increased expression of GFP under a fluorescence dissecting microscope (Zeiss Discovery V8). Complementation tests were carried out by generating worms heterozygous for two mutant alleles and scoring *pals-5p::GFP* fluorescence. Linkage group mapping with visible markers was done using the strains ERT507, ERT508, and ERT509 and scoring *pals-5p::GFP* fluorescence (see Table 1 for strain genotypes). For whole-genome sequencing analysis of mutants, genomic DNA was prepared using a Puregene Core kit (Qiagen) and 20X sequencing coverage was obtained using a 90 bp paired-end Illumina HiSeq 2000 at Beijing Genomics Institute. We identified only one gene (*pals-22*) on LGIII containing variants predicted to alter function in all three mutants sequenced (*jy1, jy2*, and *jy3*). The *jy1* mutation is an early stop at amino acid 51, *jy2* is an early stop at amino acid 105, and *jy3* is a G to A mutation of the splice acceptor before the 4th exon.

### CRISPR deletion of 11 pals genes

To construct a 34kb *C17H1* region deletion mutant, we used the co-conversion strategy as described (Arribere et al., 2014) with *dpy-10* sgRNA as the selection marker. We first constructed the short guide RNA (sgRNA) plasmids (*C17H1.3*_sg2 and *C17H1.7*_sg2) by designing the sgRNA (5’-gaacagagtgaagcaggaag-3’) to target *C17H1.3/pals-2* and sgRNA (5’-acgggcagatatacagagac-3’) to target *C17H1.7/pals-6*. The short oligonucleotides were synthesized (IDT), annealed and ligated into a modified version of DR274 (Addgene Plasmid #42250) where the sgRNA site was flanked by BsaI and *C. elegans* U6 promoter and terminator from pU6::klp-12_sgRNA (Gift from Michael Nonet). The constructed sgRNA expression plasmids (20 ng/uL each) were co-injected into N2 worms with *C17H1* region single stranded donor DNA (5’-attttgctcttatcacatttatagaaatgacaaaagtcaccgagccctcggtttttctttgcgatagttcagagcttctcaaatctctca-3’) (500nM), pDD162 (Addgene Plasmid #47549) that expresses Cas9 with empty sgRNA (50ng/uL), *dpy-10(cn64)* single stranded donor DNA (500nM), and *dpy-10(cn64)* sgRNA (20 ng/uL). Dumpy or roller F1s were selected to ensure that all CRISPR/Cas9 were expressed and *dpy-10* was successfully modified. Worms were subsequently genotyped for the deletion with single worm 3-primer PCR where a primer pair flanking the whole deletion region and a third primer in the proposed deleted region (GW503 (5’-gttagaaatgcgctgtgacgt-3’), GW504, (5’-agctcgctcagcattgttg-3’), GW515 (5’-ggaatggtactaccagtgctg-3’)). The *dpy-10* animal was crossed with N2 to obtain the final ~34kb deletion mutant strain without the dumpy phenotype. *pals* genes in the deleted region are *pals-2, pals-3, pals-4, pals-5, pals-6, pals-7, pals-8, pals-9, pals-10, pals-11*, and *pals-12*.

### Gene expression analysis

To measure endogenous mRNA expression changes, synchronized L1 worms were grown at 20°C for 48 hours on OP50 or RNAi bacteria, and then collected in TriReagent (Molecular Research Center, Inc.). RNA extraction, reverse transcription, and qRT-PCR were performed as previously described (Troemel et al., 2006). Each biological replicate was measured in duplicate and normalized to the *snb-1* control gene, which did not change upon conditions tested. The Pffafl method was used for quantifying data (Pfaffl, 2001). qRT-PCR primer sequences are available upon request.

### Osmotic stress assays

Assays were performed as described (Mazzeo et al., 2012; Solomon et al., 2004). Briefly, worms were raised under standard conditions until 24 hours post-L4 stage, then transferred to plates containing 500 mM NaCl and incubated at 20°C. For polyQ aggregation assays, worms were imaged on a fluorescence dissecting microscope and scored for aggregates at the indicated timepoints. For osmotic stress resistance assays, worms were scored for motility over a period of 10 minutes.

### Heat stress assays

Worms were grown on standard NGM plates until the L4 stage at 20°C. To assess thermotolerance, worms were shifted to 37°C for two hours. Following heat shock, plates were laid in a single layer on the bench top for 30 minutes to recover, and then moved to 20°C incubator overnight. Worms were scored for survival 24 hours after heat shock; animals not pumping or responding to touch were scored as dead. To assess response to prolonged heat stress, worms were shifted to 30°C for 24 hours and then imaged or harvested for RNA extraction. Attempts to combine RNAi and heat shock assays using worms feeding on HT115 bacteria were not successful, likely due to effects of diet on physiology.

### Size measurement

Synchronized L1 stage animals were grown on NGM plates seeded with OP50 and incubated at 20°C. The COPAS Biosort machine (Union Biometrica) was used to measure the time of flight (length) of worms at the indicated timepoints. 500–700 worms were measured for each strain at each timepoint.

### RNAi experiments

Overnight cultures of RNAi feeding clones in the HT115 bacterial strain were seeded onto RNAi plates and incubated at 25°C for 1 day. Synchronized L1 stage animals were fed RNAi for 48 hours at 20°C. All experiments with feeding RNAi used an *unc-22* positive control RNAi clone, which resulted in twitching animals in all experiments.

### Fluorescence imaging of C. elegans

Images in Figure 5 were captured with a Zeiss LSM700 confocal microscope. All other *C. elegans* images were captured with a Zeiss AxioImager.

### Lifespan

L4 stage worms were transferred to 6 cm NGM plates seeded with OP50 bacteria and incubated at 25°C. Worms were scored every day, and animals not responding to touch were scored as dead. Animals that died from internal hatching or crawled off the plate were censored. Worms were transferred to new plates every day throughout the reproductive period.

### Western blot

Synchronized young adult animals (24h after the L4 stage) were washed off NGM plates with M9 buffer, and then washed twice with M9. Worms pellets were then resuspended in SDS-PAGE sample buffer (62.5 mM Tris-HCl pH6.8, 2% SDS, 0.01% Bromophenol blue, 10% glycerol, 10 mM dithiothreitol) and boiled at 95°For 10 min. Lysates were centrifuged at 8,000 g for 5 min and then supernatants were run on a 4–20% gradient SDS-PAGE gel (Bio-Rad) and transferred to Polyvinylidene fluoride (PVDF) membrane (Bio-Rad). The blots were probed with anti-GFP diluted at 1:1000 and JLA20 anti-actin antibody (Millipore) diluted at 1:1000. The blots were treated with ECL reagent (Amersham GE Healthcare Life Sciences), and imaged on a Bio-Rad ChemiDoc.

### Bioinformatic Analysis

Amino acid sequences of 36 genes were aligned using MUSCLE (version 3.8.31) (Edgar, 2004) and trimmed with trimAl (version 1.2) (Capella-Gutiérrez et al., 2009). Bayesian Markov chain Monte Carlo inference (LG + I + G + F) was performed using BEAST (version 1.8.3) (Drummond et al., 2012). Analyses were performed with a chain length of 10 million states (sampled every 1,000 iterations) applying an uncorrelated relaxed clock (lognormal distribution) and a Yule model tree prior. Convergence and mixing was assessed with Tracer (version 1.5) and maximum clade credibility tree generated after a 25% burn-in. Posterior probability values greater than 0.5 are marked on branch labels.

### smFISH staining

A set of 48 CAL Fluor Red 610-tagged smFISH probes targeting the *cul-6* transcript was obtained from Biosearch (Stellaris^®^ FISH Probes). smFISH staining was performed as described (Ji and Van Oudenaarden, 2012). Briefly, worms were fixed in 4% paraformaldehyde, then permeabilized in 70% ethanol. smFISH probes were then hybridized overnight at 30°C. After hybridization, samples were washed 3X in 10% formamide wash buffer, then mounted on slides with Vectashield plus DAPI mounting medium. Imaging was performed using a Zeiss AxioImager M1 upright fluorescent microscope with a 63X oil immersion objective. Z-stacks of anterior intestines were acquired with a z-spacing of 1 μm, collecting signal for GFP (GFP-tagged PALS-22 protein), CAL Fluor Red 610 (*cul-6* smFISH probe), DAPI (DNA staining), and DIC. Image processing was performed using ImageJ (https://imagej.nih.gov/ij/) and smFISH spot quantification was performed using StarSearch (http://rajlab.seas.upenn.edu/StarSearch/launch.html).

## Author Contributions

Conceptualization: K.C.R. and E.R.T.; Investigation: K.C.R., T.D., J.N.S., J.P., K.C., and E.S.L.; Writing: K.C.R. and E.R.T.; Supervision: E.R.T. and D.W.; Funding Acquisition: E.R.T. and D.W.

## Acknowledgements

We thank Lakshmi Somasundaram for technical assistance, and Oliver Hobert for communicating unpublished results. Some *C. elegans* strains were provided by the Caenorhabditis Genetics Center (CGC), funded by the NIH Office of Research Infrastructure Programs P40 OD010440. We thank Oliver Hobert, Aaron Reinke, Eric Bennett, and Robert Luallen for providing helpful comments on the manuscript. J.N.S. is an IRACDA fellow and was supported by NIGMS/NIH award K12GM068524. This work was supported by NIH R21 AI115442 to D.W. and by GM114139, AG052622, and a Burroughs Wellcome Fund fellowship to E.R.T.

